# Modelling the impact of curtailing antibiotic usage in food animals on antibiotic resistance in humans

**DOI:** 10.1101/077776

**Authors:** B.A.D. van Bunnik, M.E.J. Woolhouse

## Abstract

Consumption of antibiotics in food animals is increasing worldwide and is approaching, if not already surpassing, the volume consumed by humans. It is often suggested that reducing the volume of antibiotics consumed by food animals could have a public health benefits. Although this notion is widely regarded as intuitively obvious there is a lack of robust, quantitative evidence to either support or contradict the suggestion.

As a first step towards addressing this knowledge gap, we develop a simple mathematical model for exploring the generic relationship between antibiotic consumption by food animals and levels of resistant bacterial infections in humans. We investigate the impact of restricting antibiotic consumption by animals and identify which model parameters most strongly determine that impact.

Our results suggest that, for a wide range of scenarios, curtailing the volume of antibiotics consumed by food animals has, as a stand-alone measure, little impact on the level of resistance in humans. We also find that reducing the rate of transmission of resistance from animals to humans may be more effective than an equivalent reduction in the consumption of antibiotics in food animals. Moreover, the response to any intervention is strongly determined by the rate of transmission from humans to animals, an aspect which is rarely considered.

## 1. Introduction

Heightened concern about increasing levels of antimicrobial resistance worldwide has led to renewed calls to reduce substantially the use of antibiotics in food animals[1]. The notion that such a reduction could have public health benefits arises primarily from the observation that the volume of antibiotics consumed by food animals worldwide is approaching, and may have already overtaken, the volume consumed by humans [2]. This situation is expected to worsen as the transition to intensive animal production systems continues in many regions, especially China, India and other Asian countries [3, 4].

Antibiotic consumption by food animals occurs for the purposes of herd health, prophylaxis and growth promotion. Growth promotion, often involving sub-therapeutic doses, is particularly controversial. It has been banned in European Union (EU) countries since 2005 and is the subject of a more recent voluntary ban in the USA. Currently, 51% of the OIE member countries have a complete ban on using antimicrobials as growth promoters, and a further 19% have a partial ban[5, 6]. The agricultural industry has adapted to these measures in ways such that there have been only modest net impacts on consumption of antibiotics by food animals and levels of antibiotic resistance therein, such that any consequent benefits to human health are not easily discerned [7].

A key challenge for understanding the expected impact of reducing drug usage is that the relationship between antibiotic consumption by food animals and levels of resistant bacteria in humans is complex. First, food animals are far from the only source of human exposure to antibiotic resistant bacteria: high levels of antibiotic use in hospitals, clinics and the general population are also major drivers of resistance in humans [8]. Quantifying the specific contribution of the food animal route is not straightforward and has yet to be attempted. Second, there will be many different answers to the quantification question: there are numerous combinations of different antibiotics, bacterial strains and farm animal species, each with their own dynamics, and these are likely to vary between different countries with different health care systems and agricultural production systems.

As a first step towards addressing this knowledge gap, here we develop a simple mathematical model for exploring the generic relationship between antibiotic consumption by food animals and levels of resistant bacteria in humans. Our objective is to better understand the dynamics of AMR moving between food animal and human populations and to identify which model parameters have the greatest influence on levels of resistance in humans and for which parameter combinations we expect to see the greatest impact of reducing antibiotic consumption by food animals. By using the simplest possible mathematical model as a starting point, we hope to be able to make a first step understanding this highly complex system and gain some robust and useful insights into its behaviour.

## 2. Materials and Methods

### (a) Mathematical model

Our mathematical model is intended to be as simple as possible while still capturing the non-linearities inherent in infectious agents spreading between two host populations. To achieve this we consider two variables: *R_H_*, the fraction of humans with antibiotic resistant bacteria (so a measure of the ‘level of resistance’); and *R_A_*, the fraction of food animals with antibiotic resistant bacteria. The rationale of the model is that humans or food animals can acquire antibiotic resistant bacteria from different sources:

1. Within host selection for resistant bacteria in response to direct exposure to antibiotics;
2. Direct or indirect exposure to antibiotic resistant bacteria or mobile genetic elements containing resistance determinants carried by other individuals within the same population (e.g. humans acquiring resistance from a human source);
3. As (2) but between different populations (e.g. food animals acquiring resistance from a human source).

The dynamics of *R_H_* and *R_A_* are given by the coupled ordinary differential equations:

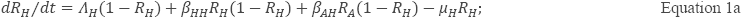

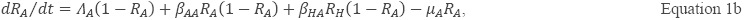

where: *Λ_H_* is the per capita rate at which humans acquire antibiotic resistant bacteria through direct exposure to antibiotics and *Λ_A_* is the equivalent in food animals; (1 − *R_H_*) and (1 – *R_A_*) are respectively the fraction of humans without antibiotic resistance and the fraction of food animals without antibiotic resistance. *β_HH_* is the per capita rate at which humans acquire antibiotic resistant bacteria as a result of exposure (directly or indirectly via environmental contamination) to other humans harbouring resistant bacteria and *β_AA_* is the equivalent for animals; *β_AH_* is the per capita rate at which humans acquire antibiotic resistant bacteria as a result of exposure (directly or, more frequently, indirectly via food products or environmental containation) to food animals harbouring resistant bacteria and *β_HA_* is the reverse; *μ_H_* is the per capita rate at which humans with resistant bacteria revert to having only susceptible bacteria (as a combination of clearance of resistance bacterial infections and demographic replacement) and *μ_A_* is the equivalent in food animals. The time unit is arbitrary and does not affect the equilibrium values. We assume that all transmission parameters (e.g. all *β's)* combine transmission of both antibiotic resistant bacteria and transmission of mobile genetic elements. Although these rates could be disaggregated, this would (for our study) have no impact on the system dynamics for a given set of parameters.

Studying the system in a steady state (i.e. at equilibrium), allows us to explore the long-term effects of changing different parameter values. To obtain an equation for the equilibrium value, 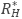, Equation 1 was set to 0 and solved for *R_H_* in terms of the eight model parameters. Although we do not consider the real-world system to be at equilibrium, largely because antibiotic consumption patterns have changed considerably in recent decades (from zero before 1932) and continue to do so, we regard 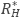 as a useful indication of where the model system is tending, and the approach to 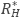 will be relatively rapid if *μ_H_* and *μ_A_* are high, i.e. the mean duration of resistant infections is short (≪1 year).

As an indication of the potential impact of curtailing antibiotic usage in farm animals on the level of resistance in the human population we define 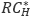 as an adjusted 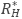 in which *Λ_A_* = 0 (i.e. no antibiotic usage in food animals) and define impact, ɷ, as 1 - the ratio of 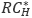 to 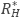:

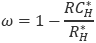

All analyses were carried out in Wolfram Mathematica, version 10.3 [9].

### (b) Scenarios

To identify key parameters and capture the non-linearities of the system in more detail we consider two distinct scenarios in this study, a low impact scenario and a high impact scenario. The difference between these two scenarios is the choice of baseline value for *β_HA_*. For the low impact scenario *β_HA_*=0.1 and for the high impact scenario *β_HA_*=0.001. These values where chosen to maximise the differences whilst minimising changes in the baseline levels of resistance between the two scenarios. The baseline parameters for the two scenarios are given in Table 1. The baseline parameter values were chosen such that the long-term prevalence of the fraction of the human population that is affected by resistant bacteria is roughly 70%. This is, for example, consistent with the situation of bacterial resistance to ampicillin in the United Kingdom, where both humans and food animals show similarly high level of resistance [10, 11].

**Table 1.** Baseline parameters for the two scenarios.

| Parameter | Description | Value used |  |
| --- | --- | --- | --- |
|  |  | Low impact | High impact |
| $\beta_{AA}$ | per capita rate at which animals acquire antibiotic resistant bacteria as a result of exposure to other animals harbouring resistant bacteria | 0.1 | 0.1 |
| $\beta_{HH}$ | per capita rate at which humans acquire antibiotic resistant bacteria as a result of exposure to other humans harbouring resistant bacteria | 0.1 | 0.1 |
| $\beta_{AH}$ | per capita rate at which humans acquire antibiotic resistant bacteria as a result of exposure to food animals carrying resistant bacteria | 0.1 | 0.1 |
| $\beta_{HA}$ | per capita rate at which food animals acquire antibiotic resistant bacteria as a result of exposure to humans carrying resistant bacteria | 0.1 | 0.001 |
| $\lambda_H$ | per capita rate at which humans acquire antibiotic resistant bacteria as a result of direct exposure to antibiotics | 0.1 | 0.1 |
| $\lambda_A$ | per capita rate at which food animals acquire antibiotic resistant bacteria as a result of direct exposure to antibiotics | 0.1 | 0.1 |
| $\mu_A$ | per capita rate at which humans with resistant bacteria revert to having only susceptible bacteria | 0.1 | 0.1 |
| $\mu_H$ | per capita rate at which food animals with resistant bacteria revert to having only susceptible bacteria | 0.1 | 0.1 |

### (c) Sensitivity analysis

We determine which model parameters have most influence on the outcome value by computing the total sensitivity index *D_Ti_* using the extension of Fourier amplitude sensitivity test (FAST) as described in Saltelli *et al.* [12]. The extended FAST method is a variance-based, global sensitivity analysis technique that has been largely used for studying complex agricultural, ecological and chemical systems (see [13, 14] for examples). Independently of any assumption about the model structure (such as linearity, monotonicity and additivity of the relationship between input factors and model output), the extended FAST method quantifies the sensitivity of the model output with respect to variations in each input parameter by means of spectral analysis. It provides measures of the amount of variance of the prevalence that arise from variations of a given parameter in what is called a total sensitivity index, *D_Ti_*. It therefore captures the overall effect of parameter variations on equilibrium levels of resistance over a pre-specified range (i.e., including first-and higher-order interactions between model parameters). For example, a value of *D_Ti_*=0.10 indicates that 10% of the total observed variation of the prevalence is explained by the parameter under consideration. The sensitivity analysis was carried out using R [15]. For the sensitivity analysis we used a parameter range of 0.0 to 1.0 for all parameters under investigation.

## 3 Results

For the two scenarios considered here Figure 1 shows the trajectory of *R_H_* and *R_A_* in time. Figure 1 shows that the long term prevalence of resistance in the human population stabilises to a value of 0.71 for the low impact scenario and 0.70 for the high impact scenario. The long term prevalence of resistance in the animal population stabilises to a value of 0.71 for the low impact scenario and 0.62 for the high impact scenario. These differences between the scenarios are due to the lower value of *β_HA_* in the high impact scenario (Table 1). These prevalences are consistent with scenarios encountered in practice (see Methods).

**Figure 1.**
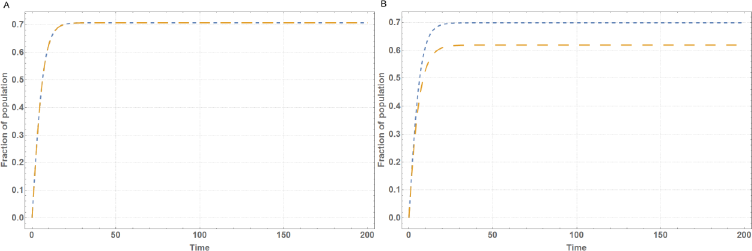
Trajectory of the fraction of the human population with antibiotic resistant bacteria (*R_H_*) and the fraction of food animals with antibiotic resistant bacteria (*R_A_*) in time for the low impact scenario (panel A) and the high impact scenario (panel B). Blue curves represent (*R_H_*), orange curves represent (*R_A_*).

The sensitivity analysis on the equilibrium equation for 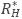 shows that this equilibrium is most sensitive to changes in *μ_H_*, followed by *Λ_H_*, *β_AH_* and *β_HH_*, (Figure 2). Furthermore, the system is minimally sensitive to changes in “animal” parameters *(β_AA_, ʲ_HA_, Λ_A_* and *μ_A_*).

**Figure 2.**
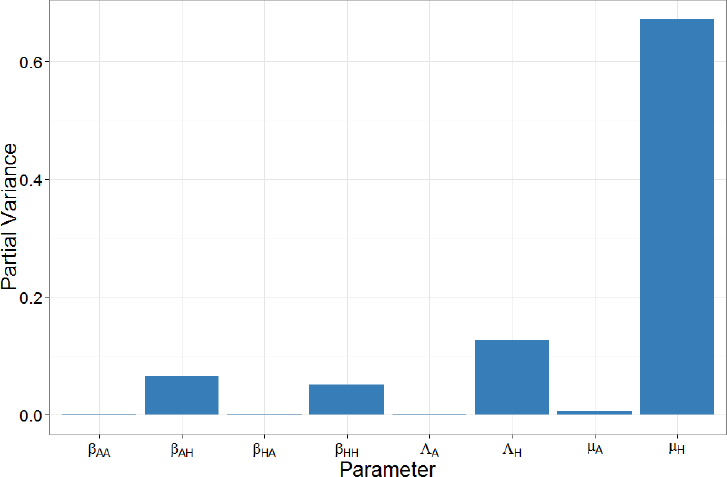
Results of a global sensitivity analysis on the equilibrium equation showing the partial variance of the individual model parameters. Higher bars indicate greater sensitivity of the model to that parameter. See Methods section for details about the sensitivity analysis and parameter ranges used.

Figure 3 shows that, in accordance with the sensitivity analysis, but covering a wider range of parameter space (0.0 to 1.0), the equilibrium 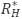 is relatively insensitive to changes Λ_A_, but is more sensitive to changes *β_AH_*, suggesting that reducing the former without addressing the latter may have limited impact on the prevalence of antibiotic resistance in the human population. Furthermore, the figure shows that the change in 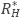 due to *β_AH_* is non-linear, which suggests that partial reductions of *β_AH_* may only have limited impact when 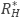 is already high. Comparing panel A with panel B in Figure 3 shows that the effect of reducing the rate at which animals acquire antibiotic resistant bacteria as a result of exposure of animals to antibiotics *(Λ_A_*) is also strongly influenced by the per capita rate at which food animals acquire antibiotic resistant bacteria as a result of exposure to humans carrying resistant bacteria *(β_HA_)*. For the higher value of *β_HA_* (low impact scenario, panel B) lowering *Λ_A_* by itself has little influence.

**Figure 3.**
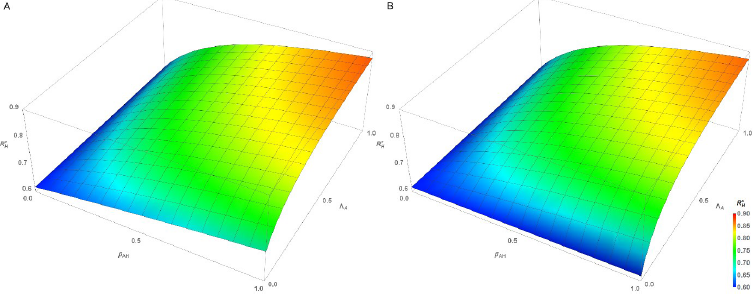
Surface of 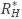 as a function of *β*_*AH*_ and *Λ*_*A*_ for the low impact scenario (panel A) and the high impact scenario (panel B).

Figures 4 and 5 show the results for the impact of curtailing antibiotic use in food animals, quantified as the variable ω (defined above). Figure 4 shows the sensitivity of the impact to the different parameters. From Figure 4 it is clear that the impact is most sensitive to *Λ_H_*, the rate at which humans acquire antibiotic resistant bacteria as a result of exposure of humans to antibiotics, followed by *μ_H_, μ_A_* and *β_HA_*. *Λ_A_* and *β_AH_* have less influence. Figure 5 shows the effects of varying *β_AH_* and *Λ_A_* on the impact, *ɷ*, of curtailing antibiotic use in food animals. The difference between the low impact scenario and the high impact scenario is immediately clear from these graphs as there is virtually no impact of reducing *Λ_A_* in the low impact scenario (with relatively high per capita rate of transmission from humans to food animals, *β_HA_*) while in the high impact scenario (with relatively low *β_HA_*) there is a much more obvious benefit.

**Figure 4.**
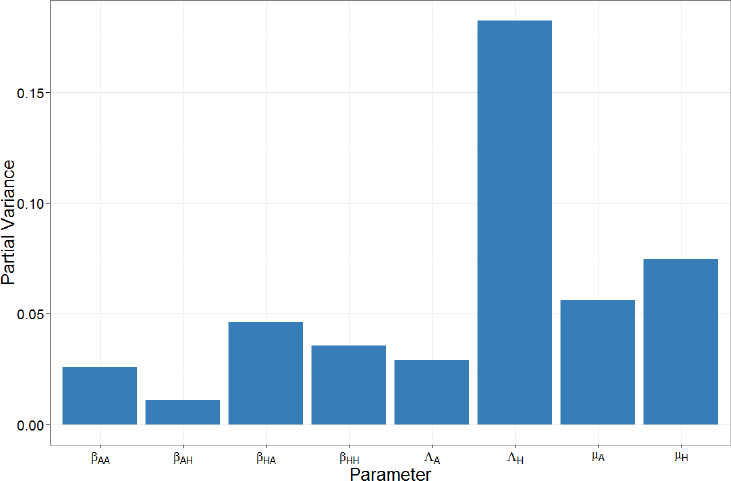
Results of a global sensitivity analysis on the impact (ω, defined in the Methods section) showing the partial variance of the individual model parameters. Higher bars indicate greater sensitivity of the model to that parameter. See Methods section for details about the sensitivity analysis and parameter ranges used.

**Figure 5.**
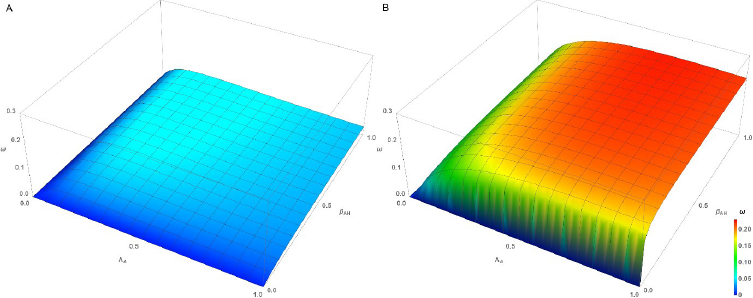
Surface of impact, ω, as a function of *β_AH_* and *Λ_A_* for the low impact scenario (panel A) and the high impact scenario (panel B).

## 4. Discussion

It is often implied that the high levels of consumption of medically important antibiotics by food animals is contributing significantly towards the global public health problem of antibiotic resistance. Therefore we tested the potential impact of curtailing the use of antibiotics in food animals on the (long term) prevalence of humans carrying resistant bacteria using a mathematical model designed to capture the non-linearities inherent in the transmission of infectious agents between two populations as simply as possible. Our results show that, as expected, the system is sensitive to changes in per capita rate at which humans acquire antibiotic resistant bacteria as a result of direct exposure to antibiotics (*Λ_H_*). Of much greater interest is the importance of the per capita rate of transmission of antimicrobial resistance from humans to animals (*β_HA_*) (see Figures 4 and 5). For this reason, we compared two scenarios, a low impact scenario (high *β_HA_*) and a high impact scenario (low *β_HA_*). If *β_HA_* is high (Figure 3, panel B; Figure 5, panel B) then the effects of reducing the rates at which animals acquire resistance as a result of antibiotic usage *(Λ_A_*) and humans acquire antibiotic resistant bacteria from animals (*β_HA_*) are limited (Figure 3 and Figure 5, panel A, when *β_AH_* or *Λ_A_* approaches 0). This contrasts with the situation where *β_HA_* is low (Figure 3, Figure 5, panel A). This indicates that whenever the rate of transmission of antibiotic resistant bacteria from humans to animals is high it is more difficult to curb the antibiotic resistance problem, a rather counterintuitive result and often overlooked in discussion about this topic.

Also of interest is that a failure to address the agricultural usage of antibiotics severely limits what can be achieved by tackling the problem from the human side, i.e. even if no resistance is required via direct exposure to antibiotics in humans (achieved by reducing *Λ_H_* to 0), we can only reduce the long term prevalence of antibiotic resistance in humans to 0.56 for the low impact scenario and 0.54 for the high impact scenario (see Figure S1 in the Supplementary information). In other words, if resistance dynamics in human and animal populations are coupled, as is generally thought to be the case in practice, substantial impacts on levels of resistance requires coordinated interventions across both populations. Our study has several limitations that should be recognised. The first limitation is the simplicity of the model used. As indicated in the introduction, antibiotic resistance is a highly complex problem with numerous routes of transmission and with dynamics that will vary qualitatively and quantitatively for different drug-bug-animal combinations. Accounting for all these different routes and combinations separately is challenging. However, by taking the simplest possible mathematical model as a starting point, we are able to make a first step in trying to understand this highly complex system and gain some robust and useful insights into its behaviour. These findings can then be used as stepping stones for the development of more complicated (and perhaps more precise) models. As with all models, several assumptions have been made in this study. For example, *Λ* is clearly related to antibiotic consumption, but the shape of this relationship is left undefined in this study as we are only interested here in the specific alternative scenario where *Λ_A_* = 0. For a partial reduction in *Λ_A_*, however, this relationship would need to be specified as different relationships could lead to (very) different results. Similarly, for the *β*'s we combined the rate of acquiring resistance through transmission of (clonal) bacteria and through the transmission of mobile genetic elements. Although it is possible to disaggregate these modes of transmission, this would require an additional parameter and would not alter the system dynamics for a given set of parameters (and thus would not alter our results). However, if the separate contributions of the different transmission paths were of interest, such disaggregation would be necessary. Lastly, for the recovery rates (μ) we assumed that these combine recovery from infection with a resistant strain with demographic replacement by hosts with susceptible ones. The scenarios considered here assume high recovery rates and so emphasize the former (given that human and food animal demographies are very different).

Key aspects of future models might include: i) information on geographical distance and contact structures between and within populations; ii) explicit consideration different modes of inheritance of resistance; and iii) quantitatively relating rates of gain of resistance to historical data on antibiotic consumption. Also, the dependency of resistance on demographics should be taken into account as human and livestock demographics can be very different (for example the batch structure that is common in poultry has an influence on infectious disease [16]).

As with all models, parameterisation is an important issue. In this study we chose our baseline parameter values such that the long term prevalence of the fraction of the human population that is affected by resistant bacteria is roughly 70%. This mimics, for example, the situation of ampicillin resistance in both human and livestock in the United Kingdom where antibiotic resistance is well established but still leaves room for improvement or deterioration of the levels of resistance. However, knowledge of the prevalence of resistance in the two populations is not by itself sufficient to parameterise this model, as there are eight parameters to be estimated. Independent estimates of parameter values are required, but are not currently available, even for any specific drug-bug-animal combination in a defined setting. In practice, the many different combinations of antibiotic, bacteria strain, food animal species and setting will represent many different points in parameter space, each of which would need to be determined individually – a significant challenge. That said, the results of the simple, generic model presented here are robust in the sense that they apply over a wide range of parameter space that we expect to cover many real world scenarios.

## 5. Conclusions

To conclude, we have shown in this study that we can obtain useful insights into a highly complex problem like antibiotic resistance by using a simple mathematical model. Although it is widely regarded as intuitively obvious that reducing antibiotic consumption in animals would decrease levels of antibiotic resistance in humans this is, in fact, not the case for a wide range of scenarios (i.e. parameter space), especially if this intervention is made in isolation. Reducing the rate of transmission of resistance from animals to humans may often be more effective. In addition, the behaviour of the system, and so the response to any intervention, is strongly determined by the rate of transmission from humans to animals, but this has received almost no attention in the literature. It is thus not enough to only lower the consumption of antibiotics in food animals, the transmission both from and to food animals should also be limited in order to maximise the impact of this and other interventions. We recommend that formal, quantitative analyses are needed to assess the expected benefits to human health of reducing antibiotic consumption by food animals. In some circumstances these benefits will be very small and other measures will be needed to reduce the public health burden of antibiotic resistance.

## Acknowledgements

We thank colleagues from Edinburgh Infectious Diseases for their helpful discussions during the drafting of this manuscript.

## Funding Statement

This study was financially supported by the *European Union's Horizon 2020 research and innovation programme* under grant agreement number 643476 (www.compare-europe.eu). The funders had no role in study design, data collection and analysis, decision to publish, or preparation of the manuscript.

## Competing Interests

We have no competing interests.

## Authors' Contributions

BvB participated in the design of the study, performed the analysis and drafted of the manuscript. MW conceived of the study, designed the study and helped draft the manuscript. All authors gave final approval for publication.

## References

1. Woolhouse M., Ward M., van Bunnik B., Farrar J. 2015 Antimicrobial resistance in humans, livestock and the wider environment. Philosophical Transactions of the Royal Society of London B: Biological Sciences 370(1670).

2. The independent Review on Antimicrobial Resistance Chaired by Jim O'Neill, (2015). Review on antimicrobial resistance. Antimicrobials in agriculture and the environment: reducing unnecessary use and waste. (Series) Review on antimicrobial resistance. Antimicrobials in agriculture and the environment: reducing unnecessary use and waste. Available at: http://amr-review.org/sites/default/files/Antimicrobials%20in%20agriculture%20and%20the%20environment%20%20Reducing%20unnecessary%20use%20and%20waste.pdf [Accessed: 12-07-2016].

3. Robinson T.P., Pozzi F. 2011 Mapping supply and demand for animal-source foods to 2030. Rome, Animal Production and Health Working Paper. No. 2.

4. Alexandratos N., Bruinsma J., (2012). World agriculture towards 2030/2050: the 2012 revision. (Series) World agriculture towards 2030/2050: the 2012 revision. Available at: [Accessed:

5. Diaz F. 2013 Antimicrobial use in animals: Analysis of the OIE survey on monitoring of the quantities of antimicrobial agents used in animals

6. Grace D. 2015 Review of evidence on antimicrobial resistance and animal agriculture in developing countries.

7. Jensen H.H., Hayes D.J. 2014 Impact of Denmark's ban on antimicrobials for growth promotion. Current Opinion in Microbiology 19, 30–36. (doi:http://dx.doi.org/10.1016/j.mib.2014.05.020).

8. Holmes A.H., Moore L.S.P., Sundsfjord A., Steinbakk M., Regmi S., Karkey A., Guerin P.J., Piddock L.J.V. Understanding the mechanisms and drivers of antimicrobial resistance. The Lancet 387(10014), 176–187. (doi:http://dx.doi.org/10.1016/S0140-6736(15)00473-0).

9. Wolfram Research, Inc,. 2015 Mathematica. 10.2 ed. Champaign, Illinois, Wolfram Research, Inc.

10. EARS-Net. 2010 European Antimicrobial Resistance Surveillance Network.

11. UK-VARSS 2014, (2014). UK Veterinary Antibiotic Resistance and Sales Surveillance Report. (Series) UK Veterinary Antibiotic Resistance and Sales Surveillance Report. Available at: https://www.gov.uk/government/uploads/system/uploads/attachmentdata/file/477788/Optimised_version_VARSS_Report_2014_Sales_Resistance_.pdf [Accessed: 18-12-2016].

12. Saltelli A., Tarantola S., Chan K.P.S. 1999 A quantitative model-independent method for global sensitivity analysis of model output. Technometrics 41(1), 39–56. (doi:10.2307/1270993).

13. Makowski D., Naud C., Jeuffroy M.-H., Barbottin A., Monod H. 2006 Global sensitivity analysis for calculating the contribution of genetic parameters to the variance of crop model prediction. Reliability Engineering & System Safety 91(10-11), 1142–1147. (doi:10.1016/j.ress.2005.11.015).

14. Neumann M.B., Gujer W., von Gunten U. 2009 Global sensitivity analysis for model-based prediction of oxidative micropollutant transformation during drinking water treatment. Water Research 43(4), 997–1004. (doi:10.1016/j.watres.2008.11.049).

15. R Core Team. 2015 R: A Language and Environment for Statistical Computing. Vienna, Austria.

16. Rozins C., Day T. 2016 Disease eradication on large industrial farms. Journal of Mathematical Biology 73(4), 885–902. (doi:10.1007/s00285-016-0973-9).

